# Integrating Multisource Target Evidence with Machine-Learning Potency Prediction for Alzheimer’s Disease Drug Discovery

**DOI:** 10.64898/2026.10.14.759026

**Authors:** Anyu Pan, Tianhua Zhai, Jonghyun Lee, Charles Zheng, Dokyoon Kim, Li Shen

**Affiliations:** University of Pennsylvania, Philadelphia, PA, USA

## Abstract

Alzheimer’s disease (AD) target evidence is distributed across biological, chemical, structural, regulatory, and trial-registry sources, complicating reproducible target evaluation and modeling. We developed a traceable data-to-model platform that integrates AGORA-centered evidence through Target Explorer, Drug-Target Lookup, and Model Portfolio workspaces. The resource contains 967 targets, 452,249 compounds, and 1,430,260 measurements. On a curated PIK3CA dataset, target-specific fine-tuning of the Boltz-2 affinity head increased Pearson r from 0.248 to 0.560 and reduced mean absolute error from 0.900 to 0.706 on 469 held-out compounds. Separately, we screened 42 regression models over 50 random splits for ten moderate-sized target datasets; a random forest model for XDH achieved the highest mean R^2^ (0.815). The platform connects curated evidence, model-ready datasets, reproducible evaluation, and public results while clearly separating contextual evidence from therapeutic claims and random-split performance from generalization to unseen chemical scaffolds.

## Introduction

Alzheimer’s disease (AD) is biologically diverse and remains a major therapeutic challenge despite recent advances in disease-modifying treatments. Its biology involves amyloid and tau, immune and vascular pathways, lipid metabolism, endosomal trafficking, and other interacting processes.^1^ Consequently, evaluating a nominated target requires more than a disease-association score. Researchers must also determine whether relevant compounds and assays are available, whether the protein identity and structure are well defined, and what is known about related drugs, regulatory records, and clinical studies.

The required evidence is distributed across resources designed for different purposes. The Accelerating Medicines Partnership–Alzheimer’s Disease (AMP-AD) AGORA platform prioritizes genes and proteins using human genetics and multi-omics evidence.^2^ BindingDB and ChEMBL curate biochemical activities and compound–target relationships,^3, 4^ whereas UniProt and the Protein Data Bank (PDB) provide protein and structure records.^5, 6^ Drugs@FDA and ClinicalTrials.gov describe regulatory products and registered studies, respectively.^7, 8^ Because these resources use different identifiers, update schedules, molecule formats, activity units, and target definitions, a combined record can hide important distinctions, such as a salt versus its parent compound, a multi-protein target, a censored activity value, or a clinical study that mentions a drug without testing the target under review.

Fragmentation also separates evidence review from predictive modeling. A researcher may identify a target in one interface, reconstruct its activity set through ad hoc exports, and train a model without preserving the source data or exclusion decisions. Model performance then becomes difficult to interpret or reproduce. This problem is especially important for datasets containing only a few hundred compounds, where strong random-split performance may reflect predictions for familiar chemical series rather than generalization to new scaffolds. Structure-aware foundation models can also be adapted to target-specific potency data, but their gains are informative only when training, validation, and held-out testing are clearly separated.

We address this gap with three contributions. **First**, we built an AGORA-centered target database and a Target Explorer for reviewing multisource evidence. **Second**, we developed a Drug-Target Lookup that resolves a named drug and AD target to pair-specific measurements, molecule identity, and FDA and ClinicalTrials.gov context. **Third**, we used the same evidence base to select modeling datasets and built a Model Portfolio containing PIK3CA-specific Boltz-2 affinity fine-tuning, downloadable model weights, and classical machine-learning baselines for ten moderate-data targets.^9, 10^ We evaluated the resulting workflow through evidence coverage and mapping summaries, held-out PIK3CA testing, and repeated-split classical-model benchmarks.

## Methods

### Study design and data sources

The workflow covered data collection, record matching, evidence integration, dataset construction, model evaluation, and public access to results. We integrated data from AGORA, BindingDB, ChEMBL, UniProt, RCSB PDB, Drugs@FDA, and ClinicalTrials.gov. Source files and API records were collected through August 11, 2026. We recorded each source, retrieval date, and processing step; all reported counts refer to this study snapshot.

### Target, protein, activity, and structure integration

AGORA targets formed the entry set. Ensembl identifiers were first mapped to all human UniProt candidates. When no Ensembl bridge was available, we allowed an exact human HGNC-symbol match. We retained all candidates and designated a preferred record using reviewed status, canonical status, and exact gene-symbol support. Original identifiers, mapping method, candidate rank, and ambiguity status remained available for review.

BindingDB and ChEMBL records were standardized while retaining activity type, value qualifier, assay description, source document, and original value. Explicit molar concentration units were converted to nM, and a derived activity scale was calculated as *p*Activity = 9 *−* log_10_(nM). Exact and censored measurements were stored separately. Multi-protein and complex targets were preserved as source entities; an activity was assigned to an individual protein only when the source relationship identified that component unambiguously.

Compound processing used RDKit-based sanitization and normalization while preserving the supplied structure and stereochemistry.^11^ Standardized full structures and fragment-parent structures were stored separately, and Bemis–Murcko scaffolds were derived to check overlap between data splits.^12^ Records that normalized to the same structure could be compared across sources without removing the original rows. UniProt mappings were joined to PDB records, structure metadata, and ligand annotations. The resulting tables were stored in a read-only DuckDB database for fast filtering and reproducible export.^13^

### Regulatory and clinical-registry context

Drug names, active ingredients, synonyms, and ChEMBL identifiers were grouped into drug families. An exact Drugs@FDA active-ingredient match to one unambiguous ChEMBL family was required; salt and solvate forms were grouped with the parent drug, whereas prodrugs remained distinct. The broad linkage set included all activity-derived target connections for the matched drug family. The high-confidence linkage set required at least one exact quantitative measurement from a direct, human single-protein assay that passed the data-quality filters. FDA application and product records, openFDA labels, and Orange or Purple Book context were kept separate from biochemical target evidence.

ClinicalTrials.gov was searched using the matched drug identity, and duplicate records were combined by National Clinical Trial (NCT) number. The interface reports registry matches separately and does not treat them as FDA approval or evidence of target-specific efficacy. Regulatory and registry records were stored in separate read-only tables so that later website updates would not change the study data used here.

### Public evidence workspaces

We developed three read-only workspaces. Target Explorer supports filtering across AGORA nomination evidence, activity availability, protein mappings, structures, and data-quality indicators. Drug-Target Lookup accepts a drug name or ChEMBL identifier together with an AGORA gene symbol and returns pair-specific measurements, assays, publications, structures, and separately labeled FDA and registry context. Model Portfolio connects selected datasets to model descriptions, evaluation results, and downloadable files where available. Each workspace provides its underlying row-level tables as CSV files and identifies the relevant data sources and limitations. The application, documentation, and available files are accessible at https://agora-evidence-ae3f8403.azurewebsites.net/.

### PIK3CA dataset construction

PIK3CA was selected because the integrated evidence contained enough quantitative IC_50_ data for separate training, validation, and testing; selection reflected data availability, not therapeutic priority. We retained exact, positive IC_50_ measurements from high-confidence, direct, human single-protein biochemical assays. Records also had to pass structure, assay, and data-quality filters, including removal of exact source duplicates, variants, mutants, covalent assays, PAINS matches, and replicate groups spanning more than one log unit. Remaining replicate values within the same parent compound, assay, and mechanism group were summarized by the median. Parents whose retained values differed by more than one log unit across assay groups were excluded. The resulting main dataset contained 4,300 observations from 4,224 unique parent compounds; 66 parents contributed more than one retained observation.

To prevent closely related records from appearing in different partitions, observations sharing a parent compound or source document were kept together. A block was eligible for the temporal held-out test only when its parent compounds, activities, and source documents were absent from an earlier ChEMBL reference snapshot and the documents entered ChEMBL after that snapshot. The remaining blocks were ordered by ChEMBL release and publication year. Newest blocks entered validation until its 15% target was reached; the rest formed training.

The split produced 3,110 observations from 3,072 parents for training, 721 from 683 for validation, and 469 observations from 469 unique parents for held-out testing. No parent compound or source document appeared in more than one partition. The training, validation, and test sets contained 1,182, 256, and 249 unique Bemis–Murcko scaffolds, respectively. Four validation scaffolds (1.6%) also occurred in training, 9 test scaffolds (3.6%) occurred in training, and 18 test scaffolds (7.2%) occurred in validation. Thus, the evaluation was separated by parent compound and source document but not by scaffold. A sensitivity analysis excluding compounds with unresolved atom stereocenters contained 3,923 observations from 3,861 compounds, including 405 held-out test observations.

The affinity-head label followed the Boltz-2 convention, *y* = log_10_(IC_50_/*µ*M), with lower values indicating greater measured potency. For visualization and the classical benchmark, observed and predicted values were converted to *p*IC_50_ = 6 *− y*. This transformation reverses the numerical direction of potency but leaves correlations and absolute-error magnitudes unchanged.

### PIK3CA model adaptation and evaluation

Using identical protein–ligand inputs, we compared the released Boltz-2 model with a model whose affinity head was fine-tuned for PIK3CA.^9, 10^ Approximately 8.7 million affinity-head parameters were trainable; the structure-prediction component remained fixed. Training used a Huber loss (*δ* = 0.5), AdamW optimization with a peak learning rate of 1 *×* 10^*−*4^ and no weight decay, a warm-up followed by stepwise learning-rate decay, bfloat16 mixed precision, an effective batch size of 16, gradient clipping, and ten epochs. The model with the highest validation-set Pearson correlation was used for held-out evaluation. Two additional training runs followed the same protocol and validation-based model selection.

Each input contained the full 1,068-residue P42336 sequence and the ligand’s canonical isomeric SMILES. The original and fine-tuned models used the same processed inputs and inference settings. All 469 held-out compounds produced valid predictions. The held-out test set was used only for final evaluation, not for training, validation, hyperparameter choice, or data-cleaning decisions.

Evaluation metrics were Pearson *r*, Spearman *ρ*, mean absolute error (MAE), and root mean squared error (RMSE). We estimated paired 95% bootstrap confidence intervals for the between-model difference in each evaluation metric using 10,000 compound-level bootstrap resamples. The 405-observation subset without unresolved stereocenters was evaluated as a sensitivity analysis. For context, we also trained two ligand-only baselines on the same split: a molecular-weight linear model and a random forest using 2,048-bit radius-2 ECFP4 fingerprints. These baselines were not used to select the fine-tuned model.

### Classical machine-learning benchmark

A preliminary screen identified 340 human single-protein targets with 160–1,000 directly linked parent compounds and eligible IC_50_ records. After full curation, 61 retained 190–310 parents and met minimum requirements of 30 scaffolds, five source documents, five assay groups, a two-log potency range, and five connected parent–scaffold–document groups; no single group could contain more than half of a target’s compounds. Before examining model performance, we selected ten by preferring datasets with 200–300 compounds, less concentration in the largest connected group, and sample sizes closest to 250. Each selected target had 225–295 curated parent compounds. Together, the ten datasets contained 2,523 target-specific compound instances (2,510 globally unique parent compounds): CSNK2A1, HDAC7, NR3C1, PARP2, PDE7A, PPARA, PRKCQ, PTGS1, TEK, and XDH. Within each target dataset, measurements were summarized as one median log_10_(IC_50_/*µ*M) value per parent compound.

For each compound, PaDEL-Descriptor generated the 881-bit PubChem structural fingerprint.^14^ We screened 42 regression models with LazyPredict using scikit-learn, LightGBM, and XGBoost. The screen covered linear, kernel, ensemble, boosting, and tree-based methods, including Random Forest, LightGBM, and XGBoost.^15–17^ Models used standard package settings and fixed randomization. Each training split was processed independently using mean imputation and standardization. Target-level model summaries and downloadable results are available in Model Portfolio.

For each target, we repeated a random 80/20 train–test split 50 times using fixed randomization. Only models that completed all 50 runs were ranked. Models were ordered within each target by mean test *R*^2^; mean Pearson *r*, mean MAE, and the standard deviation of *R*^2^ were also reported. For display, potency was expressed as *p*IC_50_ = 6 *−* log_10_(IC_50_/*µ*M).

Mean performance across the 50 splits was the primary benchmark result, and variability across splits was also reported. The Model Portfolio additionally displays the XDH Random Forest split with the highest test *R*^2^ as an illustrative predicted-versus-actual plot. This single split was selected only for visualization and was not used for target or model comparisons.

## Results

### Integrated evidence coverage

The study database connected 967 AGORA targets to 452,249 standardized compounds and 1,430,260 biochemical measurements (Table 1). Because one measurement can map to more than one component of a complex target, the database contained 1,472,169 measurement–target links. Structural integration produced 12,098 target-associated structure rows referencing 9,173 unique PDB entries. These linked tables allowed users to move from an AGORA target to individual activities, source assays and documents, standardized and original molecule representations, protein identifiers, structures, and ligands while retaining the original source relationship.

**Table 1.** Coverage of the study data snapshot. Counts describe different entities or relationship tables and are not additive.

| Component | Count | Interpretation |
| --- | --- | --- |
| AGORA targets | 967 | Target entry entities |
| Standardized compounds | 452,249 | Cross-source compound entities |
| Biochemical measurements | 1,430,260 | BindingDB (365,226) plus ChEMBL (1,065,034) |
| Measurement–target links | 1,472,169 | Includes supported component links for source complex targets |
| Target-associated structures | 12,098 | Structure rows referencing 9,173 unique PDB entries |
| FDA-matched canonical drugs | 1,079 | Exact resolved families; 1,076 linked to at least one target |
| Broad drug–target pairs | 9,787 | Activity-derived links involving 183 AGORA targets |
| High-confidence drug–target pairs | 2,248 | Direct links: 785 drugs and 133 targets |
| Unique ClinicalTrials.gov studies | 175,938 | Deduplicated registry records returned by drug-based queries |

Target-to-protein mapping results are reported separately from coverage (Table 2). Preferred UniProt records were available for 964 of 967 targets. Among the 964 mapped targets, 67 had one retained human UniProt candidate and 897 had more than one. The preferred record came from the Ensembl bridge for 962 targets and from an exact gene-symbol match for two.

**Table 2.** Target-to-UniProt mapping summary. Categories can overlap and percentages use the denominator shown.

| Mapping category | Count | Percentage | Denominator |
| --- | --- | --- | --- |
| Preferred UniProt record available | 964 | 99.7% | All 967 AGORA targets |
| No preferred mapping | 3 | 0.3% | All 967 AGORA targets |
| Preferred record from Ensembl bridge | 962 | 99.8% | 964 mapped targets |
| Exact HGNC-symbol fallback | 2 | 0.2% | 964 mapped targets |
| Single retained candidate | 67 | 7.0% | 964 mapped targets |
| Multiple retained candidates | 897 | 93.0% | 964 mapped targets |
| Reviewed canonical record with exact symbol | 957 | 99.3% | 964 mapped targets |

### Regulatory and registry linkage

Drug matching identified 1,079 FDA-matched drug families. Of these, 1,076 linked to at least one biochemical target relationship, yielding 9,787 broad drug–target pairs across 183 AGORA targets; three FDA-matched families had no supported target link. Applying the high-confidence rule retained 2,248 pairs spanning 785 drugs and 133 targets. ClinicalTrials.gov searches produced 343,226 drug–query–study links representing 175,938 unique study identifiers. These registry links are reported as contextual coverage and do not imply target-specific therapeutic efficacy.

### Public evidence navigation

Figure 1A summarizes the path from source data through record matching and model-ready datasets to evaluation and public results. Target Explorer allows users to select any AGORA target and apply evidence filters to review nomination context, biochemical activities, protein mappings, and structural information; Figure 1B shows CSF1R under one selected filter configuration. Drug-Target Lookup accepts a user-entered drug name or ChEMBL identifier together with an AGORA gene symbol and returns resolved drug identity, pair-specific biochemical evidence, and separately labeled FDA and ClinicalTrials.gov context; Figure 1C shows the results of a selegiline–MAOB query. Model Portfolio presents the PIK3CA-specific Boltz-2 evaluation and allows users to select a classical machine-learning target and regression model to examine performance summaries and predicted-versus-actual results; Figure 1D shows XDH Random Forest as one selected example. All interactive views and downloads use the same underlying tables as the reported counts.

**Figure 1.**
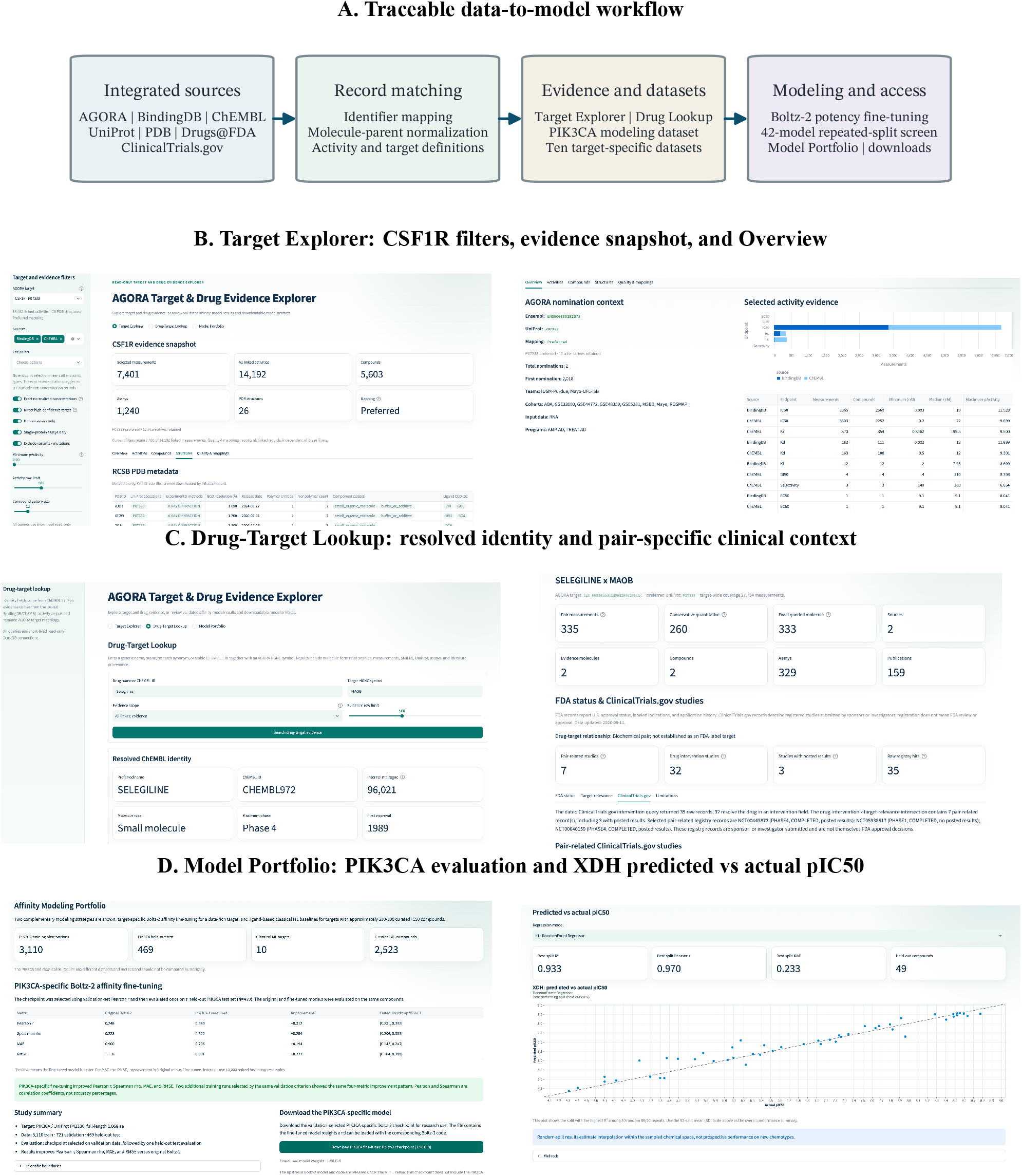
Data workflow and public application. (**A**) source data remain connected to record-matching decisions, model-ready datasets, evaluation, and downloadable files. (**B**) Target Explorer shows the selected filters, CSF1R evidence snapshot, and RCSB PDB metadata (left), with AGORA nomination context and activity summaries in Overview (right). (**C**) Drug-Target Lookup resolves the queried selegiline identity (left) before presenting the selegiline–MAOB pair summary and selected ClinicalTrials.gov records (right). (**D**) Model Portfolio presents PIK3CA held-out test results and the downloadable checkpoint (left) alongside the XDH Random Forest predicted-versus-actual pIC50 plot for the single best-performing split (right). The best-split plot is illustrative; model comparisons are based on mean performance across all 50 splits.

### PIK3CA-specific Boltz-2 fine-tuning

All four metrics improved on the held-out test of 469 unique parent compounds (Table 3). Fine-tuning increased Pearson *r* from 0.248 to 0.560 (difference 0.312; 95% CI, 0.231–0.392) and Spearman *ρ* from 0.228 to 0.522 (difference 0.294; 95% CI, 0.206–0.383). MAE decreased from 0.900 to 0.706 (improvement 0.194; 95% CI, 0.142–0.247), and RMSE decreased from 1.118 to 0.891 (improvement 0.227; 95% CI, 0.164–0.288). Both models produced predictions for all 469 compounds.

**Table 3.** Original and PIK3CA-fine-tuned Boltz-2 performance on the held-out test set (*n* = 469). Error metrics are in log_10_(IC_50_/*µ*M). Improvement is fine-tuned minus original for correlations and original minus fine-tuned for errors.

| Metric | Original | Fine-tuned | Improvement | Paired 95% CI |
| --- | --- | --- | --- | --- |
| Pearson $r$ | 0.248 | **0.560** | 0.312 | 0.231–0.392 |
| Spearman $\rho$ | 0.228 | **0.522** | 0.294 | 0.206–0.383 |
| MAE | 0.900 | **0.706** | 0.194 | 0.142–0.247 |
| RMSE | 1.118 | **0.891** | 0.227 | 0.164–0.288 |

Two additional training runs also improved every metric relative to the original model. Across all three runs, Pearson improvement ranged from 0.312 to 0.351, Spearman improvement from 0.291 to 0.352, MAE improvement from 0.194 to 0.241, and RMSE improvement from 0.227 to 0.251. In the 405-observation sensitivity analysis, the main fine-tuned model improved Pearson *r* from 0.309 to 0.573 and Spearman *ρ* from 0.292 to 0.539 while reducing MAE from 0.869 to 0.705 and RMSE from 1.091 to 0.887. The same four-metric improvement remained after compounds with unresolved atom stereocenters were excluded.

Ligand-only controls showed that compound features accounted for substantial predictive signal (Table 4). The ECFP4 Random Forest achieved Pearson *r* = 0.467 and MAE = 0.744, outperforming the original Boltz-2 model on all four metrics but remaining below the fine-tuned model. The molecular-weight linear model had the weakest performance among the evaluated methods. These controls place the fine-tuning gain in context: the adapted model achieved the best held-out performance among the evaluated methods, but the experiment does not show that the gain came specifically from three-dimensional protein–ligand information. The PIK3CA/p85 test subgroup contained only two observations and was too small for a separate analysis.

**Table 4.** PIK3CA held-out test comparison with ligand-only controls (*n* = 469). MAE and RMSE are in log_10_(IC_50_/*µ*M).

| Model | Pearson $r$ | Spearman $\rho$ | MAE | RMSE |
| --- | --- | --- | --- | --- |
| Molecular-weight linear model | 0.094 | −0.014 | 0.918 | 1.108 |
| ECFP4 Random Forest | 0.467 | 0.346 | 0.744 | 0.933 |
| Original Boltz-2 | 0.248 | 0.228 | 0.900 | 1.118 |
| PIK3CA-fine-tuned Boltz-2 | **0.560** | **0.522** | **0.706** | **0.891** |

Figure 2 complements the tables by showing held-out test predictions and the uncertainty of the improvement across the main and two additional training runs.

**Figure 2.**
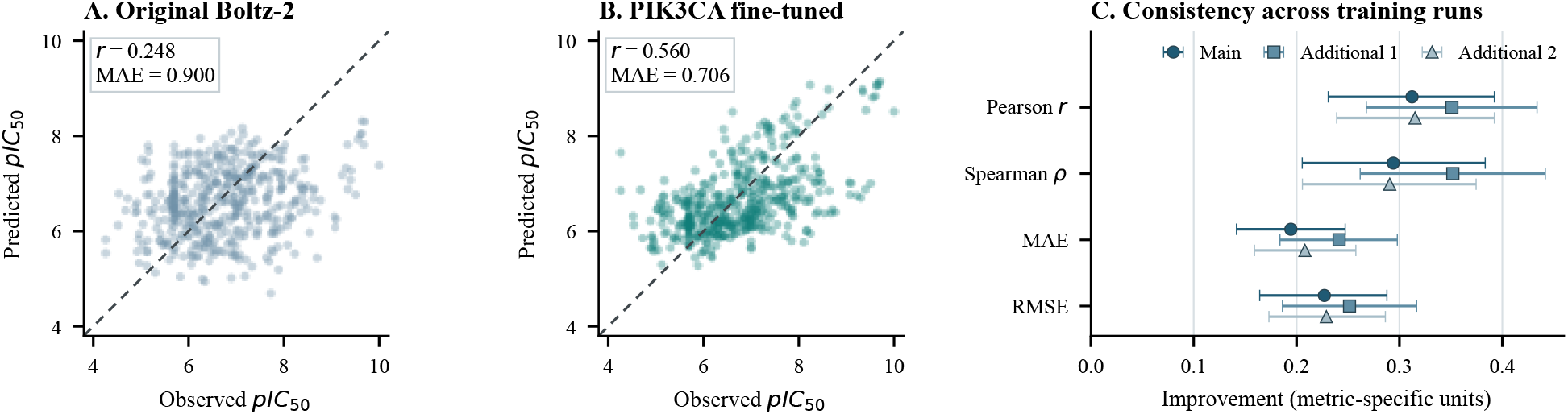
PIK3CA held-out test results for 469 unique parent compounds. **A–B**, observed versus predicted *p*IC_50_ for the original and main fine-tuned Boltz-2 models; the dashed line indicates perfect agreement. **C**, improvement in the main and two additional training runs; positive values favor fine-tuning, and whiskers show paired bootstrap 95% confidence intervals from 10,000 resamples. Correlation and error metrics use different scales and should not be compared directly.

### Ten-target classical machine-learning screen

Table 5 reports the model with the highest observed mean *R*^2^ for each target across all 50 splits. Random Forest achieved the highest mean *R*^2^ for six targets, LightGBM for two, and Gradient Boosting and Elastic Net CV for one each. A random forest for XDH had the highest mean *R*^2^ among these ten target-level summaries (0.815 *±* 0.072), with mean Pearson *r* = 0.907 and MAE = 0.332. Six other targets—PARP2, PDE7A, PTGS1, NR3C1, CSNK2A1, and PRKCQ—also achieved mean *R*^2^ *>* 0.50 for their highest-mean models. In contrast, HDAC7 and TEK had mean *R*^2^ *<* 0.40, illustrating that comparable sample counts did not guarantee comparable predictability.

**Table 5.** Model with the highest observed mean test *R*^2^ for each target across 50 random 80/20 splits. Pearson *r* and MAE are split means; MAE is on the *p*IC_50_ scale.

| Target | $n$ | Highest-mean model | $R^2$, mean (SD) | Pearson $r$ | MAE |
| --- | --- | --- | --- | --- | --- |
| CSNK2A1 | 295 | Random Forest | 0.548 (0.082) | 0.752 | 0.545 |
| HDAC7 | 228 | Random Forest | 0.388 (0.125) | 0.640 | 0.610 |
| NR3C1 | 225 | LightGBM | 0.562 (0.099) | 0.762 | 0.598 |
| PARP2 | 262 | Random Forest | 0.724 (0.084) | 0.858 | 0.526 |
| PDE7A | 234 | Random Forest | 0.703 (0.105) | 0.844 | 0.445 |
| PPARA | 235 | LightGBM | 0.479 (0.136) | 0.711 | 0.466 |
| PRKCQ | 278 | Gradient Boosting | 0.511 (0.093) | 0.732 | 0.603 |
| PTGS1 | 230 | Random Forest | 0.591 (0.118) | 0.778 | 0.383 |
| TEK | 293 | Elastic Net CV | 0.397 (0.099) | 0.645 | 0.703 |
| XDH | 243 | Random Forest | **0.815 (0.072)** | **0.907** | **0.332** |

Thirteen models ranked among the top five for at least one target, and their ten-target matrix shows clear differences among datasets (Figure 3A). Random Forest ranked in the top five for nine targets and had the highest mean *R*^2^ for six. Figure 3B shows the mean and between-split standard deviation for each target’s highest-mean model. Figure 3C displays all 50 split-level test *R*^2^ values for the XDH Random Forest (mean, 0.815; median, 0.827; SD, 0.072).

**Figure 3.**
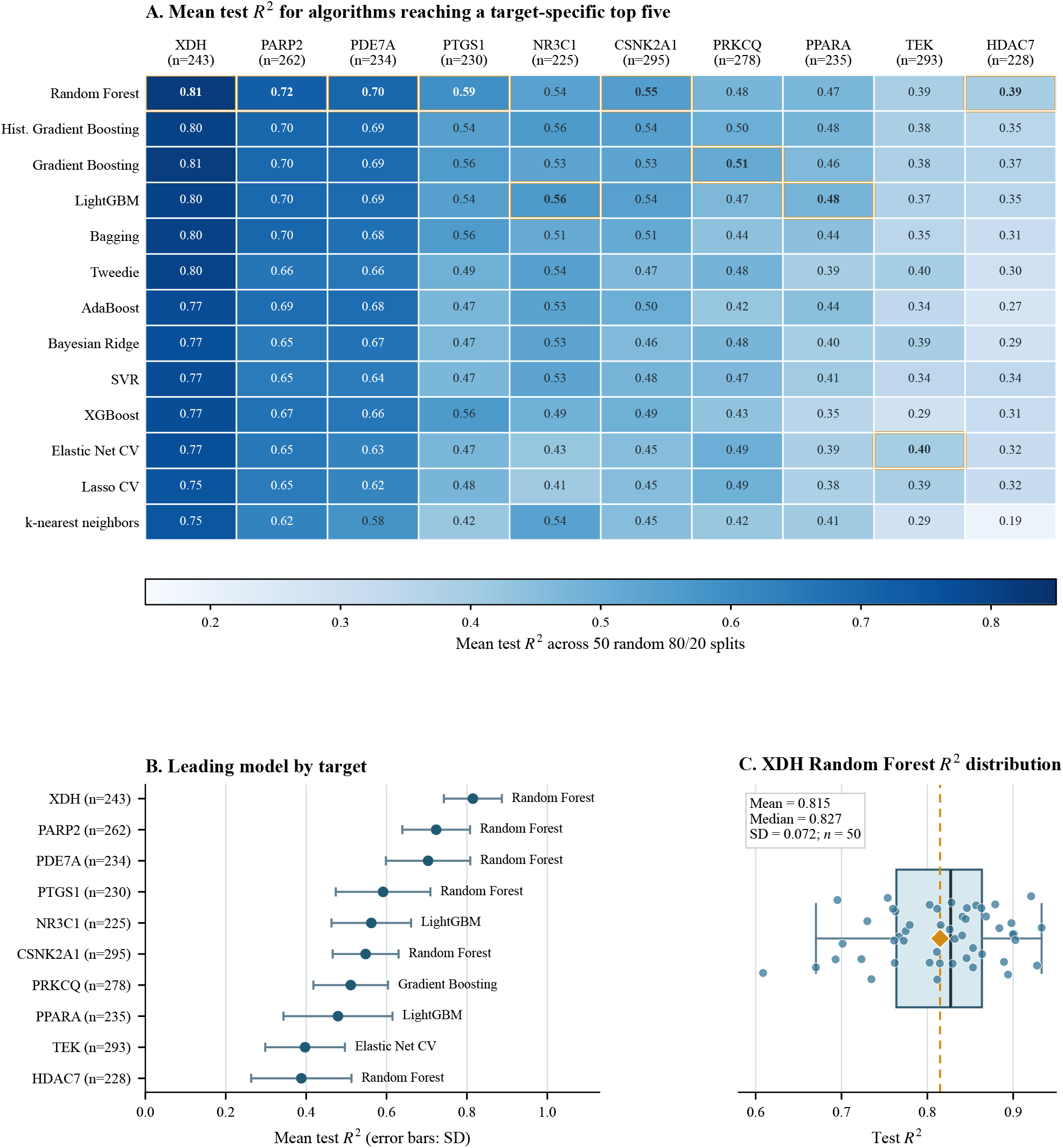
Ten-target classical machine-learning overview. All 42 regression models were screened. **A**, mean test *R*^2^ for the 13 models that ranked among the top five for at least one target and completed all 50 splits; outlined cells identify each target’s highest-mean model. **B**, mean *R*^2^ and between-split SD for each target’s highest-mean model. **C**, distribution of all 50 XDH Random Forest test *R*^2^ values; points represent individual splits, the box spans the interquartile range with the median shown by its center line, whiskers extend to 1.5 times the interquartile range, and the diamond marks the mean.

## Discussion

The three public interfaces form one data-to-model workflow rather than three separate projects. Target Explorer supports target-level evidence review, Drug-Target Lookup provides pair-specific biochemical and clinical context, and Model Portfolio connects selected datasets to model evaluation and downloadable files. Because the interfaces use the same underlying tables, users can trace a model result back to the activities, assays, documents, protein mappings, and compound representations used to construct its dataset.

Keeping evidence types separate was essential because biochemical activities, FDA product records, and clinical-trial records answer different questions. Seeing these records together can aid navigation and hypothesis formation, but it does not show that a registered study evaluated the queried target or that a drug is effective for AD. Similarly, an activity measured for a protein complex should not automatically be assigned to every component, and a matched salt form should remain distinguishable from the submitted molecule. The platform therefore shows labeled relationships and downloadable source rows rather than combining all evidence into one score.

The PIK3CA experiment illustrates the value of prespecified data partitioning and validation-only model selection. Model selection used validation results, followed by one evaluation on the held-out test set. The main model and two additional training runs all improved the four evaluation metrics on the same 469 compounds. The paired bootstrap confidence intervals did not include zero, and the sensitivity analysis showed the same direction of improvement. These results support a focused conclusion: affinity-head fine-tuning improved potency prediction on this held-out dataset relative to the original model.

These results do not establish a biological mechanism or therapeutic benefit. The ECFP4 Random Forest performed strongly, showing that ligand features alone captured substantial predictive signal. Although the fine-tuned Boltz-2 model achieved the best evaluated result, this comparison cannot determine whether the additional gain came specifically from three-dimensional protein–ligand modeling. The model also omitted the p85 regulatory chain, exact experimental constructs were not always available, some scaffolds occurred in more than one partition, and the PIK3CA/p85 test subset was too small for a separate analysis. Pose accuracy, pocket accuracy, prospective potency, selectivity, and AD-related biological effects were not evaluated.

The classical screen serves a different purpose. Repeated random splits show average performance and split-to-split variation for molecular fingerprints and commonly used regression models. The 50-split averages and distributions, rather than the single XDH interface example in Figure 1D, provide the basis for comparison. Because random splitting can place related chemical series in both training and test sets, these estimates are useful for selecting baseline models and checking the workflow, but they do not measure performance on entirely new chemical scaffolds. Differences among targets with similar sample sizes also suggest that activity noise, chemical diversity, target biology, and assay composition may matter as much as the number of compounds.

Several broader limitations remain. The data were collected through August 11, 2026, and later source updates are not represented. Automated record matching can miss valid links or create incorrect ones despite the quality checks.

Source assays use different protocols, protein constructs, endpoints, and reporting practices that cannot be removed fully by unit and structure standardization. The modeling examples cover one fine-tuned structure-aware target and ten target datasets rather than all 967 targets. We also did not have external laboratory measurements or prospective medicinal-chemistry decisions for validation. Public access improves transparency but does not replace independent replication.

Future work will prioritize external and later-collected test sets, validation on unseen scaffolds or chemical series, better uncertainty estimates, and clearer reporting of where each model is expected to work. Structure-aware models should also be evaluated with complete biological assemblies and experiment-specific protein constructs when these data are available. For evidence integration, documented updates and expert review can focus on uncertain record matches. These extensions would preserve the platform’s central principle: model claims should remain connected to the evidence and evaluation conditions that produced them.

## Conclusion

We developed a public, traceable platform that connects 967 AGORA targets, 452,249 standardized compounds, and 1,430,260 biochemical measurements through Target Explorer, Drug-Target Lookup, and Model Portfolio. Together, these workspaces link evidence review and drug–target context to model-ready datasets and public results. PIK3CA-specific fine-tuning of the Boltz-2 affinity head improved four metrics on a held-out test set, and repeated-split benchmarking provided classical ligand-based baselines for ten target datasets. The results do not establish therapeutic efficacy, a three-dimensional mechanism, or performance on new chemical scaffolds, but they provide a clear foundation for external and prospective validation.

## Acknowledgements

This work was supported in part by NIH Grants R01 AG071470, U01 AG068057, and U01 AG066833.

